# CRF receptor type 1 modulates the nigrostriatal dopamine projection and facilitates cognitive flexibility after acute and chronic stress

**DOI:** 10.1101/2022.10.26.513963

**Authors:** S Becchi, CL Burton, M Tsoukalas, J Bowring, BW Balleine, D Mor

## Abstract

Chronic unpredictable stress (CUS) impairs cognitive flexibility in rats, particularly when faced with additional mild acute stress (AS). We tested the hypothesis that this impairment is associated with alterations in dopamine activity in the dorsal striatum driven by corticotropin-releasing-factor receptor type 1 (CRFR1) in the substantia nigra pars compacta (SNpc). In experiment 1, rats received CUS or handling for 14 days, before learning two action-outcome associations (lever presses and food rewards). Learning was assessed using outcome devaluation. Cognitive flexibility was then assessed by reversing the outcome identities followed by a second outcome devaluation test, with half of the rats in each group receiving AS prior to reversal training. Dopamine and its metabolite were quantified in the dorsal striatum and CRFR1 mRNA was quantified in the SNpc. Increased dopaminergic activity in the left dorsal striatum and CRFR1 expression in the left SNpc were associated with resilience to AS in naïve rats but with impairment in CUS+AS rats, suggesting a transition in hemispheric control from left to right as a protective mechanism following CUS. This suggestion was tested in experiment 2, where SNpc CRFR1 was blocked unilaterally prior to AS and reversal training. Blocking CRFR1 in the left medial SNpc impaired cognitive flexibility following AS in naïve rats but restored it in CUS rats. Blocking CRFR1 in the left, but not right, lateral SNpc also impaired cognitive flexibility following AS in naïve rats but had no effect in CUS rats.

## Introduction

The dorsomedial striatum (DMS) is essential for encoding and retrieving the specific action-outcome (A-O) associations necessary for goal-directed action, both when they are initially encountered [1–3] and when subsequently updated [4]. This learning and updating depend on the modulatory influence of local dopamine (DA) release in the cortico-striatal pathway [5, 6]. Stress can alter dopamine neurotransmission in the striatum in both DMS and in the dorsolateral striatum (DLS) and alter the ability of animals to adapt flexibly when A-O associations change [7–9]. This has been shown in the mesolimbic area [8] and is accompanied by shifts in the lateralization of monoaminergic activity generally [10–13].

Different types of stress differentially affect activity in the dorsal striatum. In rodents, mild acute stress (AS) increased markers of dopaminergic activity in DMS and DLS but only in the left hemisphere in rats showing intact cognitive flexibility [14]. In contrast, chronic unpredictable stress (CUS) increases resting dopamine sensitivity in the substantia nigra *pars compacta* (SNpc) and reduces the response to rewarding stimuli, causing maladaptive striatal dopaminergic regulation [9]. One of the first neuropeptides released in response to stress is corticotrophin-releasing factor (CRF), which initiates humoral corticosterone release and acts centrally as a neuroregulator of cellular activity [15]. Its receptor, CRF-receptor 1 (CRFR1), is expressed on dopamine neurons in the SNpc [16, 17] that project ipsilaterally and topographically to the dorsal striatum. It is possible therefore that CRF input to the SNpc after CUS and AS differentially affects dopamine activity in DMS and DLS resulting in lateralised changes in DA activity and in cognitive flexibility.

To address this claim, in the current study we first conducted a major reassessment of brain chemistry in our prior study [8] to assess changes in striatal dopamine in regions of the dorsal striatum. This reanalysis found a lateralized effect of CRFR1 activation in the SNpc suggesting that CRFR1 activity in the left SNpc facilitated acquisition of new contingencies after AS in the Naïve group but impaired it when rats received CUS (i.e., CUS+AS group). To confirm this observation in experiment 2 we assessed the effect of infusing the CRFR1 antagonist Antalarmin, into either the medial or the lateral SNpc in the left or the right of Naive and CUS rats prior to AS and a test of behavioral flexibility.

## Materials and Methods

### Subjects

This experiment received ethics approval by the University of New South Wales Animal Ethics Committee (AEC number 19/64B and 20/69B). The procedures adhere to the ARRIVE guidelines [18]. Outbred Long Evans rats were housed in groups of 2-4 in cages controlled for ventilation and humidity on a 12-hour light/dark cycle with *ad-libitum* access to standard chow and water until behavioral training was initiated. A total of 126 were considered for the study: 62 rats in experiment 1 and 64 in experiment 2.

### Experiment 1. Effect of CUS and/or AS on CRFR1 expression in the SNpc, DA activity in the dorsal striatum and the consequences for behavioral flexibility

#### Apparatus

Behavioural training was conducted using 16 operant conditioning chambers (MED Associates) each enclosed within a sound and light attenuating shell. Each chamber contained a pellet dispenser that delivered grain pellets (45mg, Bioserv Biotechnologies) and a pump that delivered 20% sucrose solution (0.2ml). Chambers contained two retractable levers and a recessed magazine cantered between them. An infra-red photobeam was positioned at the threshold of the magazine to record entries. Each chamber contained a light (3W, 24V) illuminated for the duration of all behavioural sessions. Sessions were pre-programmed and controlled by microcomputers running MED Associates proprietary software (Med-PC). Lever pressing, magazine entry, reinforcer delivery as well as the presentation time of each lever was captured for each session using this software.

#### Chronic unpredictable stress

Rats were randomly assigned to Naïve or Chronic Unpredictable Stress (CUS) experimental groups. The CUS treatment involved daily restraint for two hours inside plexiglass tubes (20cm length, 6.35cm diameter, Ibisci, USA) over 14 days. Each day, the restraint was given at a different time between 6am to 10 pm to ensure the stress was unpredictable. Rats allocated to the naïve treatment were handled for a few minutes daily over the 14 days. Percentage bodyweight change was calculated at two-day intervals during the stressor period.

#### Instrumental training

Two days prior to commencing training rats were food restricted with daily intake was restricted to 15g of chow, which was maintained for the remainder of the experiment. Weight was monitored thrice weekly to ensure it remained above 85% of baseline bodyweight. Behavioural training started with one session of magazine training. Rats received 20 pellet and 20 sucrose outcomes at 15 second intervals for 15 minutes. Rats were then trained to press the levers such that, for each rat, one lever was assigned to deliver pellets and the other a sucrose solution, counterbalanced to control for any lever position preference. Each training session was divided into four periods, two on each lever in alternation. During each period, one lever was extended until either 20 outcomes were delivered or 15 minutes had elapsed after which the lever was retracted and a 2-min break was instituted after which the other lever was inserted and so on. Two training sessions were conducted each day with the order of lever presentation counterbalanced across sessions. Outcomes were delivered on a continuous reinforcement schedule for three sessions, then on a random ratio (RR)-5 schedule, i.e, the outcome was delivered after five presses on average, then three sessions on RR10 and two sessions on RR20.

#### Outcome devaluation by satiety

After this training rats completed two specific satiety-induced outcome devaluation tests on successive days. To induce specific satiety, rats were placed into clean devaluation boxes and given unrestricted access to either sucrose solution or pellets for 45 mins. They were then transferred to their operant chamber to compete a 10-minute extinction test. Both levers were extended simultaneously, however, no outcomes were delivered during the test. The lever associated with the outcome that was pre-satiated was considered the devalued lever and the other lever valued. The following day, rats received the same protocol only with the alternative outcome presented during the pre-feeding phase.

#### Mild acute stress and outcome-identity reversal

To assess the rats’ ability to update changes in action-outcome (A-O) contingencies they next received three training sessions with the outcome identities reversed; e.g., if, initially, pressing the left lever delivered a pellet and the right lever sucrose solution then pressing the left lever now delivered sucrose solution and the right lever a pellet. In this phase rats received only one training session per day structured as previously described with the outcomes delivered on an RR-10 schedule.

During this phase each of the previously generated stress groups was subdivided into two sub-groups: one received a mild acute stress (AS), involving 5 minutes force swim test, prior to each reversal training session, whereas the other received no treatment. This resulted in four groups: naïve (n=16), naïve with AS (n=13), CUS (n=17) and CUS with AS (n=16). For the forced swim test, rats were placed individually into a white plastic oval bin (100cm high, 30cm maximum diameter) filled to a height of 45cm with clear, fresh water (at 25± 1°C). The sub-groups under each stress condition were matched for performance in the first devaluation test to ensure similar levels of performance during initial training between the sub-groups. After reversal training, rats completed a second round of outcome devaluation tests conducted as described above except that, after the first test, rats received a fourth refresher RR10 training session with or without the acute stress (as appropriate).

#### Novel mild stress and euthanasia

To assess the impact of acute stress on dopaminergic markers in the dorsal striatum and corticotropin-releasing hormone receptor 1 (CRFR1) expression in the SNpc, all AS rats received a series of foot shocks before euthanasia. Rats were placed into an unfamiliar operant chamber where the stainless-steel rod floor was connected to a shock generator (Med-Associates, USA). In a single session, rats received three foot-shocks (5 mA) at random intervals over the course of 10 minutes. Rats were then transferred by hand into a novel cage where they were left for 20 minutes. Euthanasia was completed by rapid decapitation without anaesthesia. Brains were removed and the tissue block containing the midbrain was snap frozen over dry ice for in-situ hybridisation analysis. The remaining rats were similarly euthanised without any additional stress treatment.

#### In-situ hybridisation

Frozen, unfixed tissue was sliced using a cryostat (Leica Microsystems) and mounted on Superfrost Plus slides. Ten series of 14μm coronal sections were taken from −4.7 to −5.8mm from bregma. Slides were stored in slide racks at −20°C for one hour and then stored at −80°C. Identification of CRFR1 mRNA was achieved using the RNAscope® 2.5HD brown reagent kit (Advanced Cell Diagnostics, USA) and CRFR1 probe (Rn-Crhr1-C3, Advanced Cell Diagnostics, USA). Pictures showing either the left or the right SNpc were taken using the 40x objectives. CRFR1 mRNA positive profiles in each of the stress groups were counted in the left and right, medial and lateral SNpc at four levels (−4.9mm, −5.16mm, −5.4mm and −5.64mm from bregma) and presented as a total combined value.

#### Quantification of dopaminergic markers in the dorsal striatum using High-Performance Liquid Chromatography (HPLC)

Dopamine (DA) and dihydroxyphenylacetic acid (DOPAC) were quantified in the left or right, DMA or DLS, of all stress groups using HPLC.

Tissue blocks containing the left or right, DMA or DLS, were micro-dissected under a surgical microscope. Blocks were sonicated in homogenization buffer (150mM phosphoric acid and 500μM diethylene triamine pentacetic acid) and centrifuged for 25min at 16,000rpm. An aliquot from each supernatant was used to quantify total protein using the BCA Protein Assay Kit (ThermoFisher Scientific). The rest of the supernatant was filtered using an amicon-ultra© centrifuge filter with a cut-off of 3KD filter (Millipore). HPLC was performed using a Prominence system (Shimadzu, Kyoto, Japan) composed of degasser (DGU-20A3), liquid chromatographer (LC-20AD), autosampler (SIL-20A) and communications module (CBM-20A). Isocratic mobile phase contained 13% methanol, 87% 0.01 M monobasic sodium phosphate, 0.1 mM ethylenediaminetetraacetic acid, 0.65 mM 1-octane sulfonic acid, 0.5 mM triethylamine at pH 2.81, adjusted with hydrochloric acid. 7 μL samples were injected onto a Gemini C18 column (150×4.60 mm, 110 Å, 5 μm particle size; Phenomenex, Torrance, CA, USA) connected to an electrochemical detector (Antec Leyden Intro, Zoeterwoude, NV, Netherlands) with an Ag/AgCl reference electrode at a potential of +0.7 V and 35°C with a 1.5 mL/min flow rate. External standards for DA, DOPAC and HVA were run daily to produce a six-point calibration curve. Peak area analysis was done using Shimadzu CLASS-VP lab solutions version 6.11 data acquisition software. Concentrations in samples were normalised to the total protein in the tissue block.

### Experiment 2: The effect of blocking CRFR1 in the SNpc on behavioral flexibility following CUS and/or AS

Following the 14 days of CUS or daily handling, all rats underwent surgery for placement of a guide cannula 1mm above either the left or right medial, left or right lateral SNpc. Rats were anaesthetized with isofluorane (5% induction, 2% maintenance in 100% oxygen) and positioned in a stereotaxic frame (Stoelting, Wood Dale, IL, USA). They received a subcutaneous injection of Benacillin (Ilium) and Bupivacaine (Hospira) at the surgical site. An incision was made along the midline and a small hole was drilled into the skull above the target region. A microinjection guide cannula was implanted unilaterally into the lateral SNpc (See supplements material for coordinates). Four additional screws were fitted caudally, rostrally and laterally to the guide cannula drill with dental cement. Rats were given 7 days of recovery and monitored daily. To sustain CUS exposure during recovery, CUS rats received novel stress manipulations on days 4 (two hours of wet bedding), 5 (removal of water bottles overnight) and 6 (2 hours of 45-degree tilting of the home cage) of recovery.

Coordinates for guide cannula implantation: female rats: −5.25 caudal to bregma, 2.5 laterally from the midline, −6.6 below the skull); (male rats: −5.3mm caudal to bregma, 2.6mm laterally from the midline, −7mm below the skull), or medial SNpc (female rats: −5.35mm caudal to bregma, 1.25mm laterally from the midline, −7.0mm below the skull); (male rats: −5.4mm caudal to bregma, 1.3mm laterally from the midline, −7.4mm below the skull).

Rats with cannula placement outside the target region were excluded from the analysis for a final of 64 total rats for experiment 2.

### Statistical analysis

Experiments were subjected to two-way ANOVA or repeated measure two-way ANOVA, followed by Bonferroni tests for multiple comparisons when interactions or main effects were found significant. Behavioural data for both initial contingency and reversed contingency tests were also analysed using the contrast analysis in PSY software (UNSW) which provides a more fine-grained analysis of complex experimental designs. When an interaction was found, simple effect analysis was evaluated using PSY software. Pearson correlation coefficient analysis was used to reveal correlations. Devaluation factor was calculated as valued – devalued lever presses. A value of p<0.05 was considered statistically significant. Graphs originated using Graph Pad Prism 8. All other data were analysed using Prism, Version 8.0 (Graph Pad software).

## Results

### Effect of AS, CUS or their combination on behavioral flexibility

Our previous study (Mor et al 2022) used a two-part protocol to test the effect of a CUS and/or AS on behavioural flexibility, first assessing the encoding of the initial A-O identities using an outcome devaluation assessment before progressing to identity reversal with half of the naïve and CUS cohort also received an AS prior to reversal training

All behavioural analyses, including pressing rates during original and reversal contingencies and pressing after original and reversal contingencies devaluation tests and weight loss and circulating corticosterone levels following exposure to CUS, are reported in [8]. **Figure 1A** illustrates the experimental design; **Figure 1B** shows that CUS alone didn’t produce any alteration in initial A-O learning; whereas **Figure 1C** shows that the combination of CUS and AS impaired the ability of rats to update the A-O associations after outcome identity reversal.

**Figure 1.**
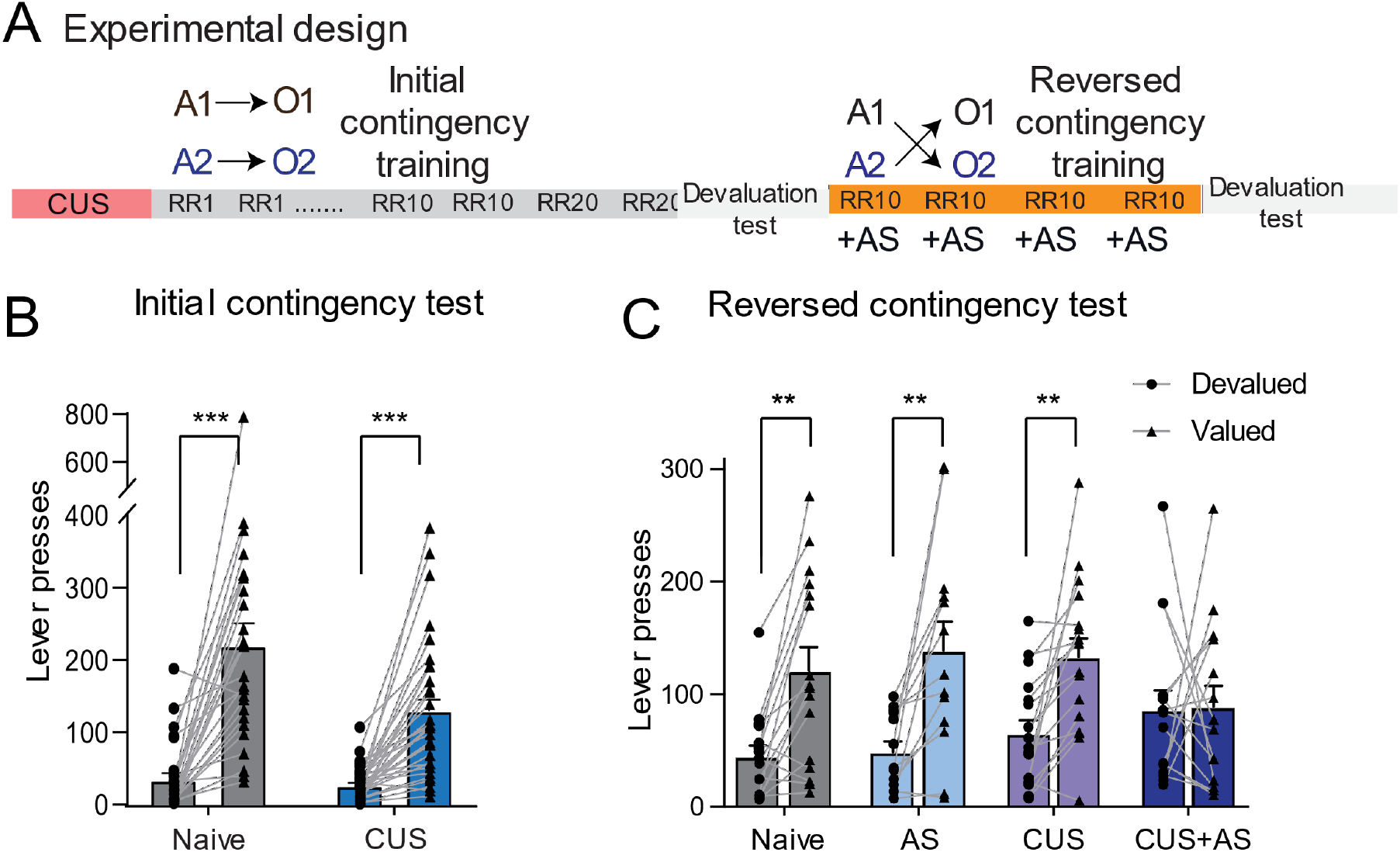
Contingency Learning Following Chronic and Acute Stress Exposure (adapted from Mor et al., 2022) (A) Design of Mor et al 2022. (B) CUS alone didn’t produce any alteration in the encoding of the initial A-O associations using an outcome devaluation test. Two-way ANOVA showed a significant devaluation effect *F*_(1,57)_=73.75, with both naïve and CUS groups pressing the value lever more than the devalued lever (*p*<.001) and no group x devaluation interaction. (C) Rats were then trained on the reversed contingency, swapping the A-O associations (see Supplementary Figure 1). In the second devaluation test, following contingency reversal, a 2×4 mixed ANOVA also revealed a significant overall effect of outcome devaluation, *F*_(1,55)_=27.81, *p*<.001 and a devaluation x group interaction F_(3,55)_=3.10, *p*=.034. Bonferroni adjusted pairwise comparisons revealed a significant devaluation effect in the *Naïve* (*p*<.001), *Naïve with AS* (*p*<.001) and *CUS* (*p*<.003) groups but not in the *CUS with AS* group (*p*=.901) indicating that the combination of CUS and AS impaired the ability to encode the changes in the A-O contingencies. Error bars represent SEM. \**p*<.05, \**p*<.01, \*\**p*<.001, *** *p*<.0001.

### Effect of AS, CUS or their combination on DA and DOPAC levels in dorsal striatum

Changes in DA and 3,4-Dihydroxyphenylacetic acid (DOPAC) levels in the left and right, DMS and DLS, following either AS, CUS or both, were assessed at the end of the behavioural task using HPLC (**Figure 2**). A detailed summary of all interactions, main effects and post-hoc analyses is available in **Table S1 in Supplementary Materials**.

**Figure 2.**
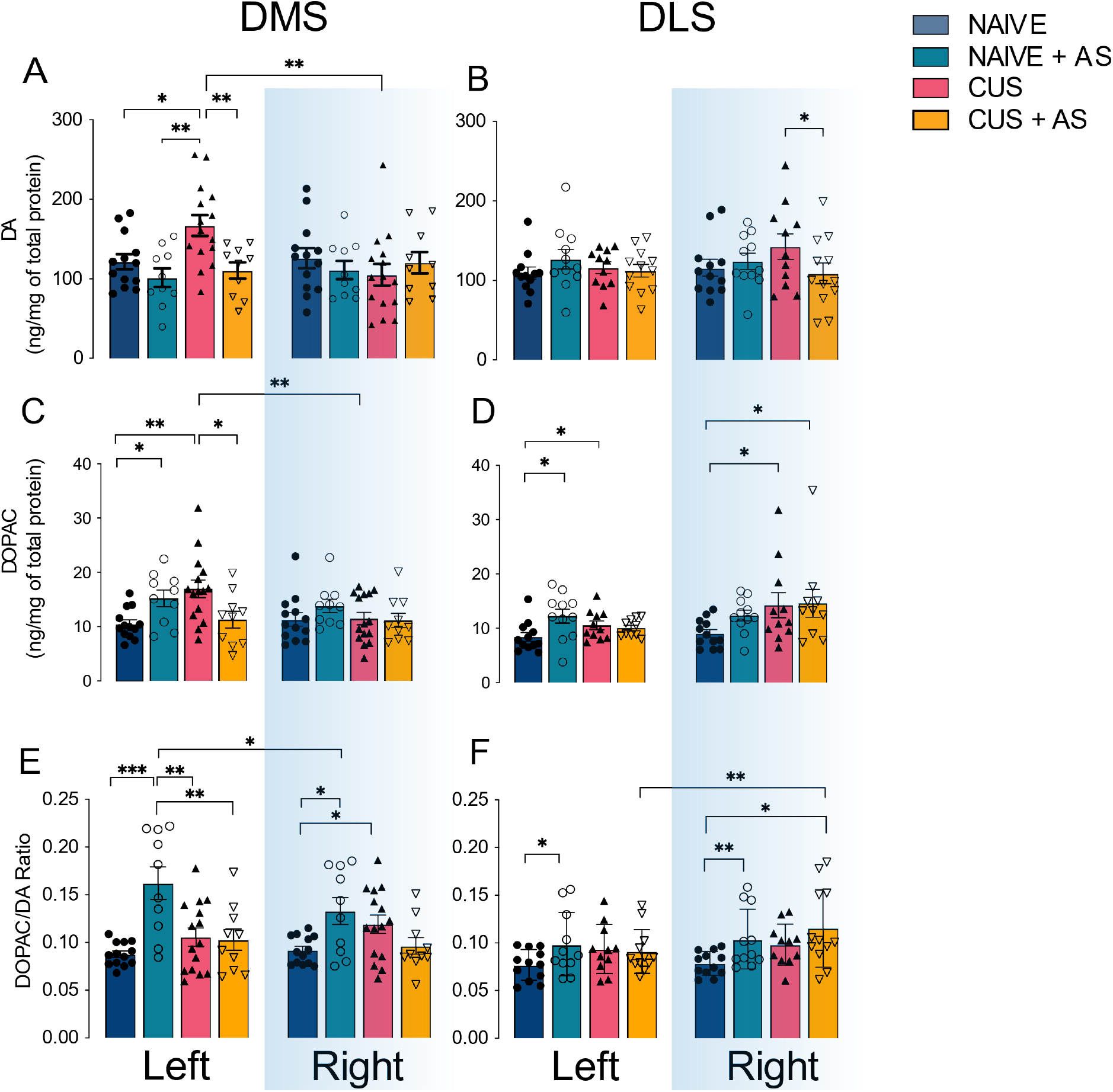
The effect of AS, CUS or the combination of both on dopamine and DOPAC in the dorsal striatum. **A,C,E**) Quantification of DA (A), DOPAC (C) and DOPAC/DA ratio (E) in the left and right hemispheres in the DMS of rats that received 2 weeks of CUS or handling with or without AS for 3 consecutive days just before reversal training. **B,D,F**) Quantification of DA (B), DOPAC (D) and DOPAC/DA ratio (F) in the left and right hemispheres in the DLS of the same rats. Bars represent means ±SEM. Naïve n=16, Naïve + AS n=13, CUS n=17, CUS+AS n=16. *\*p*< .05, \**p*< .01, \*\**p*< .001, \*\*\**p*< .0001

In the DMS, AS had differential effects on DA and DOPAC levels depending both on CUS treatment and hemisphere, resulting in three-way AS x CUS x hemisphere interactions for DA: F_(1,44)_=4.090, *p*.049 and DOPAC: F_(1,44)_=5.173,*p*=.028, and a AS x CUS interaction for the DOPAC/DA ratios F_(1,41)_=20.534, *p*<.001 (**Figure 2A, C, E**). **Figure 2A** shows increased DA after CUS compared to Naïve (*p*=.004) and a decrease to baseline level after exposure to both stressors (*p*=.001). This was evident only in the left hemisphere; no differences were found in the right hemisphere. **Figure 2C** shows that both AS and CUS alone increased DOPAC levels (*p*<.05, *p*=.001), but the combination of stressors reversed this increase to Naïve levels (*p*=.007). Again, this effect was only evident in the left hemisphere. **Figure 2E** shows that levels of DOPAC/DA ratio, which are indicative of DA turnover, increased bilaterally after exposure to AS alone (left: *p*<.001, right: *p*=.001), whereas this effect was absent in CUS. AS also led to higher DOPAC/DA ratios in the left compared to right DMS (*p*=.01).

In the DLS, we found CUS x AS interactions for DA, F_(1,40)_=8.124, *p*=.007 and DOPAC F_(1,40)_=14.942, *p*<.001.. **Figure 2B** shows that AS decreased DA in CUS rats in the right DLS, leading to significantly lower DA levels in CUS+AS when compared to CUS alone (*p*=.01). **Figure 2D** shows that AS alone increased DOPAC levels (*p*=.008) in the left and CUS alone increased DOPAC levels bilaterally (left *p*=.023, right *p*=.004). When looking at DOPAC/DA ratio in the DLS (**Figure 2F**), AS increased the ratio bilaterally (left *p*=.045, right *p*=.005).

We then looked at correlations between DA and DOPAC levels and DOPAC/DA ratios with lever-pressing rates during reversal learning sessions (S**upplementary Figure 2**) and performance at test after reversal training (**Figure 3**). **Figure 3** shows the significant and most relevant correlations. When the animals received an AS before training, the ability of the rat to encode the new A-O associations (represented as a devaluation factor, which is calculated as presses on the valued lever - presses on the devalued lever) showed a positive correlation with the DOPAC/DA ratios (r^2^=.5, *p*=.022) in the left DMS (**Figure 3A**), while a negative correlation with DA levels was found in right DLS (r^2^= .411, *p*=.024) (**Figure 3B**) and in the left DMS (r^2^=.433, *p*=.038) (**Figure 3C**). Conversely, negative correlations between devaluation factor and dopaminergic activity were found in the CUS+AS group, in both the left DMS and left DLS: DOPAC levels in the left DMS (r^2^=.697, *p*=.025) (**Figure 3D**) and DOPAC/DA in the left DLS (r^2^=.308, *p*=.049) (**Figure 3E**).

**Figure 3.**
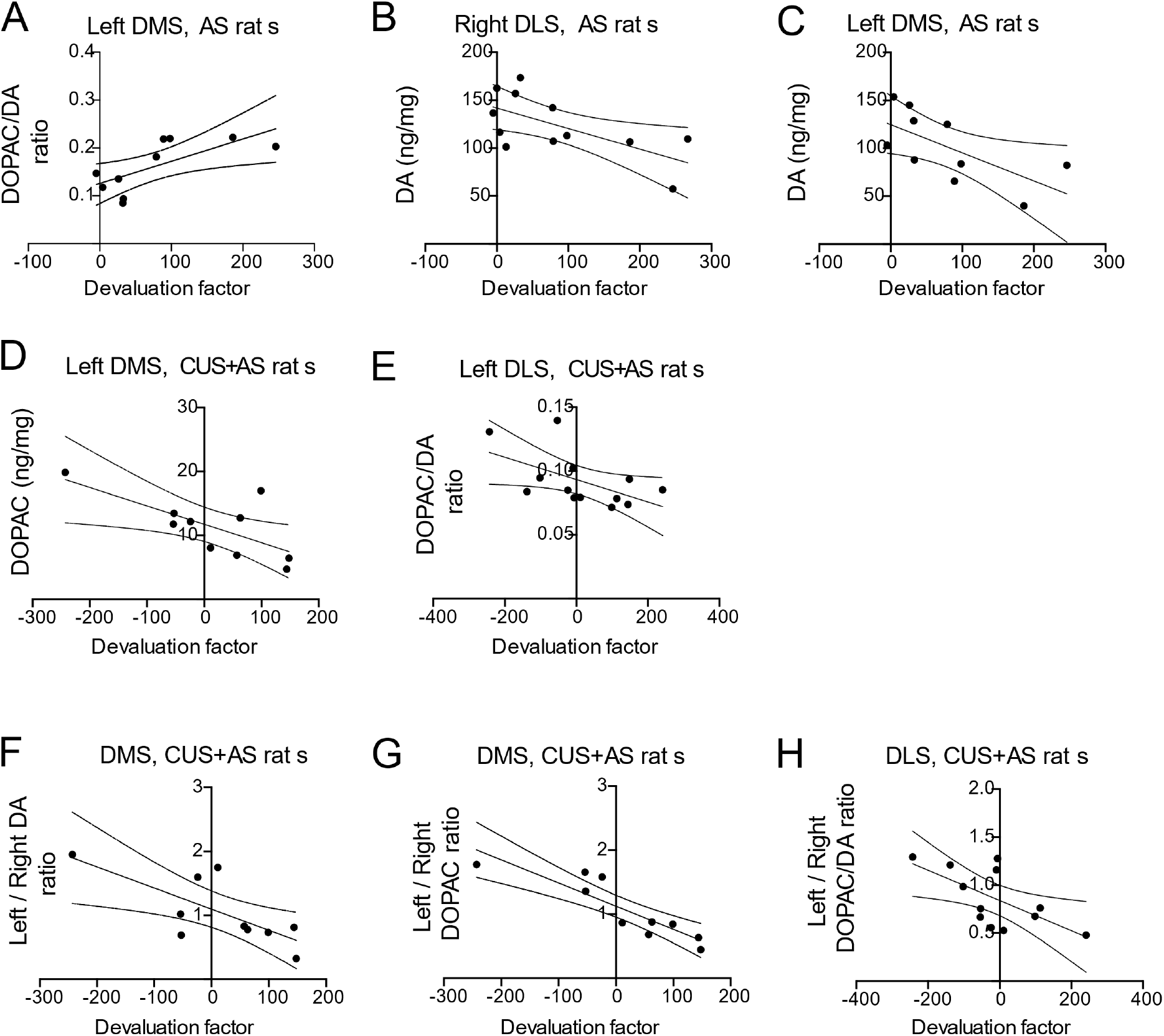
Correlations between monoamines levels and behavioral performance at test. **A**) Pearson correlation between DOPAC/DA ratios in the DMS of the left hemisphere and devaluation factor for AS group. Note that the Devaluation Factor was calculated by the presses on the value level-presses on the devalue lever, **B**) Correlation between DA level in the DLS of the right hemisphere and devaluation factor for AS rats. **C**) Correlation between DA level in the DMS of the left hemisphere and devaluation factor for AS rats. **D**) Correlation between DOPAC levels in the left DMS and devaluation factor in CUS+AS rats. **E**) Correlation between DOPAC/DA ratios in the left DLS and devaluation factor for CUS+AS rats. **F**) Correlation between the imbalance of DA between left and right hemisphere in the DMS and behavioral performance in CUS+AS rats. **G**) Correlation between the imbalance of DOPAC between left and right hemisphere in the DMS and behavioral performance in CUS+AS rats. **H**) Correlation between the imbalance of DOPAC/DA ratio between left and right hemisphere in the DLS and behavioral performance in CUS+AS rats.

Finally, to test whether the magnitude of the asymmetry between the two hemispheres played a role in updating A-O associations, correlations were performed between performance at test, after training on the reversed A-O associations, and the ratio of the monoamine levels in the left and right hemispheres of the DMS or DLS. Significant correlations were found in the CUS+AS group only and indicated reduced devaluation factor associated with higher left-to-right imbalance for DA (r^2^= .528,*p*=.017) (**Figure 3F**), DOPAC (r^2^=.787, *p*<.001) (**Figure 3G**) in the DMS and DOPAC/DA in the DLS (r^2^=.417, *p*=.023) (**Figure 3H**).

### The effect of AS, CUS or their combination on CRFR1 mRNA expression in the SNpc

CRFR1 mRNA expression in the left and right, medial and lateral SNpc was quantified using *in-situ* hybridization. Staining allowed the identification of individual mRNA molecules and a cluster of mRNA was considered a positively stained cell **(see Figure 4A)**.

**Figure 4.**
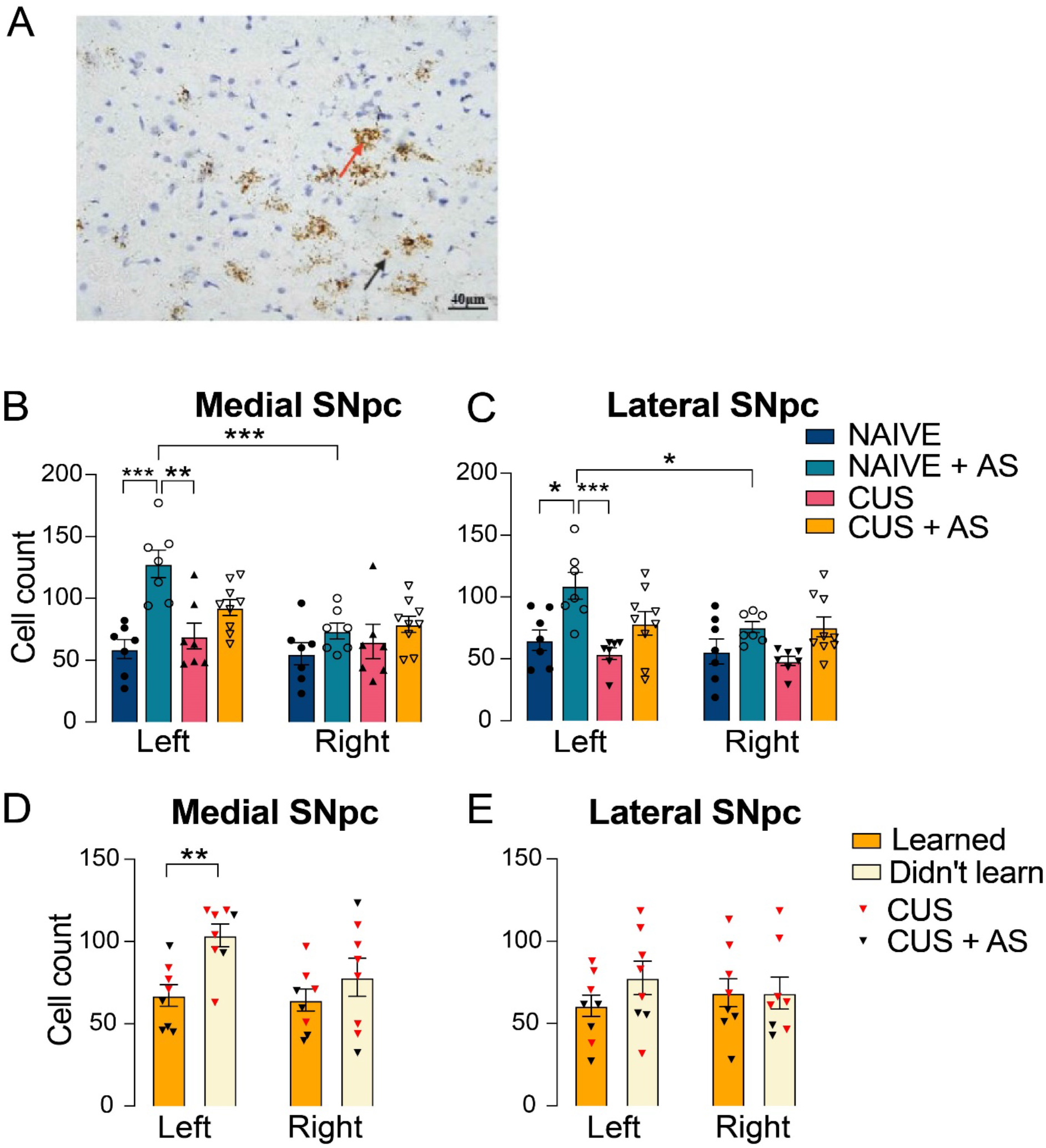
Quantification of CRFR1 in the SNpc with *in-situ* hybridization. **A**) Micrographs representing the staining of CRFR1-positive cells using *in-situ* hybridization. Staining revealed cluster of mRNAs (red arrow) or single mRNA (black arrow). Clusters of signals were counted and reported for both the medial SNpc (**B**) and the lateral SNpc (**C**) for all four groups in both left and right hemispheres. **D,E**) CRFR1-positive cell count in the medial and lateral SNpc for only CUS and CUS+AS rats, divided into two subgroups: rats that did learn the new contingency and the ones that didn’t learn it. Bars represent means ±SEM. \**p*< .05, \**p*< .01, \*\**p*< .001, *** *p*< .0001.

In the medial SNpc, a 2×4 Mixed ANOVA revealed that AS increased CRFR1-positive cells in the left medial SNpc of Naïve rats (*p*<.001), but not in CUS groups, resulting in a side x AS x CUS three-way interaction *F*_(1,26)_=6.674, *p*=.015 and a side x AS interaction *F*_(1,26)_=11.15, *p*=.002. Main effects were found for side *F*_(1,26)_=21.47, *p*<.001 and AS *F*_(1,26)_=17.67, *p*<.001 **(Figure 4B).** A similar effect was found in the lateral SNpc: a main effect was found for side *F*_(1,26)_=7.407, *p*=.011 and AS *F*_(1,26)_=18.97, *p*<.001, but not for CUS or for any of the interactions. Bonferroni multiple comparison analysis revealed that AS increased the number of immunoreactive profiles in left of both the lateral and medial SNpc but not in the right SNpc.

As shown in figure 1, the cohort of rats exposed to CUS (CUS and CUS+AS groups) showed various levels of performance, with some rats pressing more on the devalued lever and some on the valued lever, resulting in similar average performance on both levers in the CUS+AS group. We divided each group in two subgroups of rats: one that performed according to the new contingency (Learning) and one that responded according to the old contingency (No Learning). We then tested whether there was a relationship between SNpc CRFR1 expression within members of these two subgroups. A 2×2 ANOVA (learning x side) revealed a main effect for learning in the medial SNpc *F*_(1,14)_=7.55, *p*=.016, with rats that didn’t learn the reversed contingencies showing a significantly higher CRFR1-positive profiles in the left medial SNpc (Bonferroni multiple comparison, *p*= .048) **(Figure 4D)**. No differences between these learning subgroups were found in the lateral SNpc **(Figure 4E)**.

### Experiment 2: The effect of lateralized CRFR1-antagonist infusions on cognitive flexibility

HPLC and *in-situ* data from experiment 1 suggested a lateralized effect of CRFR1 activation in the SNpc on the ability to update changes in A-O associations following AS. The data suggest that CRFR1 activity in the left SNpc facilitated acquisition of new contingencies after AS in the Naïve group, but impaired it when rats received CUS (i.e., CUS+AS group). To test this hypothesis, we sought to block CRFR1 activity by infusing the antagonist Antalarmin into the medial or the lateral SNpc in either the left or the right hemisphere of rats in the Naive and CUS groups prior to the AS and reversal training.

Rats received 14 days of CUS or daily handling, during which the Naïve group gained significantly more bodyweight than CUS group (**Supplementary Figure 3A and B**), after which all rats were food-restricted and given training on the two-lever two-outcome task as previously (**Supplementary Figure 3C and D**).

A-O encoding was confirmed using sensory-specific satiety-induced outcome devaluation test. Three-way ANOVA revealed a significant devaluation effect in the lateral cannula cohort: *F*_(1, 27)_=73.885, *p*<.001, and in the medial cannula cohort: *F*_(1,29)_=57.86, *p*<.001, with no significant interactions in either cohort. Bonferroni adjusted pairwise comparisons indicated all groups encoded and retrieved A-O associations during training showing a significant difference between valued and devalued levers (**Figure 5A and C**).

**Figure 5.**
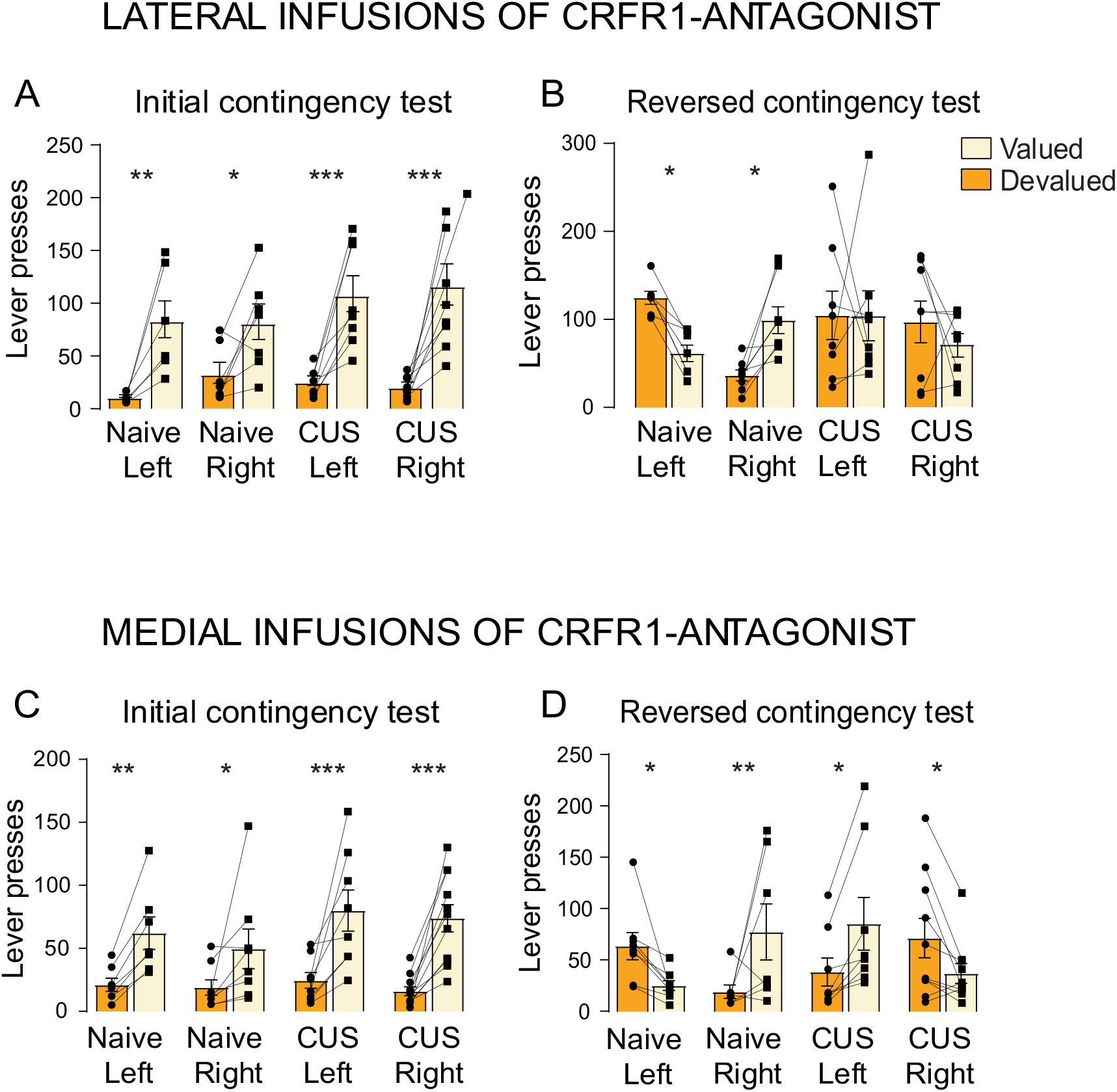
The effect of lateralized CRFR1-antagonist infusions on cognitive flexibility. **A**) Lever presses during outcome devaluation test following initial A-O association training in rats that received unilateral cannula implantation in either left or right lateral SN. **B**) Lever presses during outcome devaluation test following reversed A-O association training in rats that received unilateral infusion of CRFR1 antagonist in either left or right lateral SN just before each training session. **C**) Lever presses during outcome devaluation test following initial A-O association training in rats that received unilateral cannula implantation in either left or right medial SN. **D**) Lever presses during outcome devaluation test following reversed A-O association training in rats that received unilateral infusion of CRFR1 antagonist in either left or right medial SN just before each training session. Lateral cohort: Naïve Left n=7, Naïve Right n=8, CUS Left n=8, CUS Right n=8. Medial cohort: Naïve Left n=8, Naïve Right n=7, CUS Left n=8, CUS Right n=10. Bars represent means ±SEM. \**p*< .05, \**p*< .01, \*\**p*< .001, *** *p*< .0001.

Next, all rats received reversal contingencies training, together with an infusion of Antalarmin and AS before training (**Supplementary Figure 3)**. The second devaluation test given after reversal training revealed a three-way interaction of devaluation x CUS x side for the lateral SNpc infusion groups: *F*_(1,27)_=6.991, *p*=.014. Follow-up contrast analysis revealed an interaction between groups Naïve Right and Naïve Left x devaluation *F*_(1,27)_=9.409, *p*=.005; however no effect was observed when comparing CUS Right and CUS Left x devaluation *F*_(1,27)_<1. When comparing Naïve Left with CUS Left x devaluation, no effect was observed *F*_(1,27)_=2.321, *p*>.05, whereas an interaction was found when infusing CRFR1 antagonist in the right hemisphere, Naïve Right *vs* CUS Right x devaluation *F*_(1,27)_=4.966, *p*=.035. Simple effects analysis showed a difference between devalued and valued lever in Naïve Left *F*_(1,27)_=4.440, *p*=.045, and in Naïve Right *F*_(1,27)_=5.007, *p*=.003 but not in either CUS Left or Right groups *F*s<1 (**Figure 5B**).

A three-way devaluation x CUS x side interaction was also found in the cohort that received CRFR1 infusion in the medial SNpc: *F*_(1,29)_=26.71, *p*<.001. Follow-up contrast analysis revealed a side x devaluation effect in both Naïve and CUS: Naïve Left vs Naïve Right x devaluation *F*_(1,29)_=14.516, *p*<.001; CUS Left vs CUS Right x devaluation *F*_(1,29)_=12.194, *p*=.002; Naïve Right vs CUS Right x devaluation *F*_(1,29)_=12.138, *p*=.002; and Naïve Left vs CUS Left x devaluation *F*_(1,29)_=14.652, *p*<.001. Simple effects revealed a clear difference between devalued and valued lever in all 4 groups: Naïve Left x devaluation *F*_(1,29)_=4.922, *p*=.0345, Naïve Right x devaluation *F*_(1,29)_=9.871, *p*=.004; CUS Left x devaluation *F*_(1,29)_=7.336, *p*=.011; CUS Right x devaluation *F*_(1,29)_=4.884, *p*=.035 (**Figure 5D**).

## Discussion

We have previously shown that either a mild AS or CUS alone do not impair reversal contingencies learning, but that their combination does. Here we extend these findings by showing that intact learning following either of these stresses is associated with increased DA activity in the left DMS and in the bilateral DLS. In contrast, AS in rats previously exposed to CUS caused learning impairment and produced the opposite effect on DA activity, particularly in the left DMS and DLS. Correlation analysis indicated that differences in DA activity in the left and right hemispheres of the dorsal striatum are related to goal-directed learning and performance, with opposite hemispheric relationships observed in CUS+AS compared to Naïve and CUS rats. The *in-situ* analysis of CRFR1 found that increases in DA activity in the left hemisphere following AS were associated with increased CRFR1 expression in the medial SNpc. Although no significant differences in CRFR1 mRNA expression were found in rats exposed to CUS or combined CUS+AS, a closer analysis of these effects based on their functional influence on learning found the increased CRFR1 expression in the left medial SNpc was associated with learning impairments. There were, therefore, opposing relationships between learning and CRFR1 expression in the medial SNpc in AS and CUS+AS rats.

### Asymmetries in dorsal striatal DA activity induced by AS in naïve and CUS rats

Endogenous asymmetry in dopaminergic activity within the DS has been found in humans and rodents [11]. In non-stressful conditions, this asymmetry is associated with motor functions such as spatial preferences, direction of rotation or pressing a lever on a particular side of an instrumental apparatus [8, 19, 20], and can be presented as either left or the right dominance, depending on which limb is used or the direction of the motor movement. The functional significance of the asymmetry is not limited to motor movement; an asymmetrical increase in DA associated with directional movement also facilitates the learning associated with that action [21]. Following stress, we have found different patterns of asymmetry in DA activity in the dorsal striatum following the three different types of stress used. Intact learning following AS in naïve rats was associated with increased dopaminergic activity in the left DMS and DLS. This relationship was further supported by correlation analysis showing a positive relationship between learning and dopamine metabolism in the left DMS. This is consistent with other studies showing preferential changes in dopaminergic activity following mild stress in the left DS [14] or left cortical regions associated with the behavioral selection circuitry [10] as well as increased c-fos preferentially in the left DS following a forced swim test.

Following CUS, rats still presented with increased DA and DOPAC in the left DMS, but in the DLS, these increases were larger on the right. These were associated with reduced, but still intact learning. Finally, when CUS rats faced an additional AS, increased DA activity in the right DLS remained, whereas levels in the left DMS returned to baseline. These patterns suggest a transition from left to right with increasing severity of stress. In fact, rats with CUS+AS showed the opposite correlations between behavior and DA activity compared to AS rats, with improved learning associated with reduced activity in left DMS and DLS. Furthermore, uniquely to CUS+AS rats, the magnitude of the asymmetry, rather than the concentration in each side individually, played a role in their capacity to retain learning after AS. Rats with lower left vs right ratios in both the DMS and DLS showed better performance. Whether these associations between learning and DA asymmetry reflect asymmetries in output from striatal regions and in hemispheric activity generally, however, remains an open question.

Caution should also be exercised in interpreting changes in DA levels, as most of the DA quantified likely reflects intracellular, presynaptic storage, while levels of DOPAC or DOPAC/DA indicate increased metabolism of DA and therefore increased DA activity. An increase in total DA level could reflect increased activity in SNpc projections or local synthesis, but not necessarily increased release and utilization, something that could explain the opposite patterns found for DA and DOPAC/DA and performance in the reversal devaluation test following AS in the left DMS.

### Asymmetries in SNpc CRFR1 expression induced by AS in naïve and CUS rats

CRF modulation of the midbrain dopaminergic responses to stress has focused almost exclusively on the ventral tegmental area (VTA), even though the dopaminergic neurons of the SNpc also expressed CRFR1 [22] and have substantial CRF input from several key regions regulating behavioral responses to stress [15]. AS in naïve rats led to a significant increase in CRFR1 expression, particularly in the left medial SNpc, whereas a significant effect was not observed in CUS+AS. Similar to the patterns found for the DA increases in the left DMS, this pattern of CRFR1 expression appeared to be related to cognitive flexibility, particularly in the differences between AS and CUS+AS rats. Nevertheless, it is not clear whether activation of CRFR1 produces these effects directly. To examine this issue, in experiment 2, we blocked CRFR1 during AS prior to reversal training. We found that blocking CRFR1 in the left medial and lateral SNpc of AS rats impaired learning during the reversed A-O associations training with rats performing according to the original contingencies in the reversal devaluation test. Blocking the receptor in the right medial or lateral SNpc did not impair the learning and rats performed according to the reversed associations, further supporting the findings from experiment 1, in which CRFR1-induced dopaminergic activity in both the left DMS and DLS is required for cognitive flexibility following stress.

In contrast, blockade of CRFR1 in the left medial SNc of CUS+AS rats appeared to mitigate the learning impairment induced by combining these stress treatments. Findings from experiment 1 indicated that individual differences in learning following CUS+AS are highly dependent on the extent of the left-to-right transition in dopaminergic-driven activity. In that regard, blocking the receptor unilaterally likely polarized the CUS+AS rats into the two ends of the natural spectrum. Blocking the receptor on the left created a right dominance asymmetry and promoted learning while blocking the receptor on the right created a left dominance asymmetry and impaired learning. This effect appeared to be specific to the medial SNpc. Blocking the receptor in the lateral SNpc of CUS+AS rats, either on the left or the right neither restored nor further impaired learning. So while regression analysis in experiment 1 indicated that a transition from left to right asymmetry in the DLS is also protective in CUS+AS rats, this transition does not appear to have been mediated by CRFR1 in the lateral SNpc.

In Summary, this study further develops our understanding of the mechanisms regulating cognitive flexibility following stress and the role CRFR1 in the SNpc plays in this process. We have shown that learning is enhanced by CRFR1-driven increases in dopaminergic activity in the left dorsal striatum following AS, but impaired in CUS+AS rats. We have also shown that a left-to-right transition in the dopaminergic activity in the dorsal striatum following a combination of CUS and AS, partly driven by the opposite effect that CRFR1 in the SNpc, has a protective effect on learning.

## Author Contributions

DM: Conceptualization and design of the work

SB, DM, JB, CB, MT: Acquisition, analysis, and interpretation of data

DM, SB, JB, CB: Drafting the work

DM, BWB: Revising the work and final approval of the version to be published

## Supplementary figures and table

**Supplementary figure 1.**
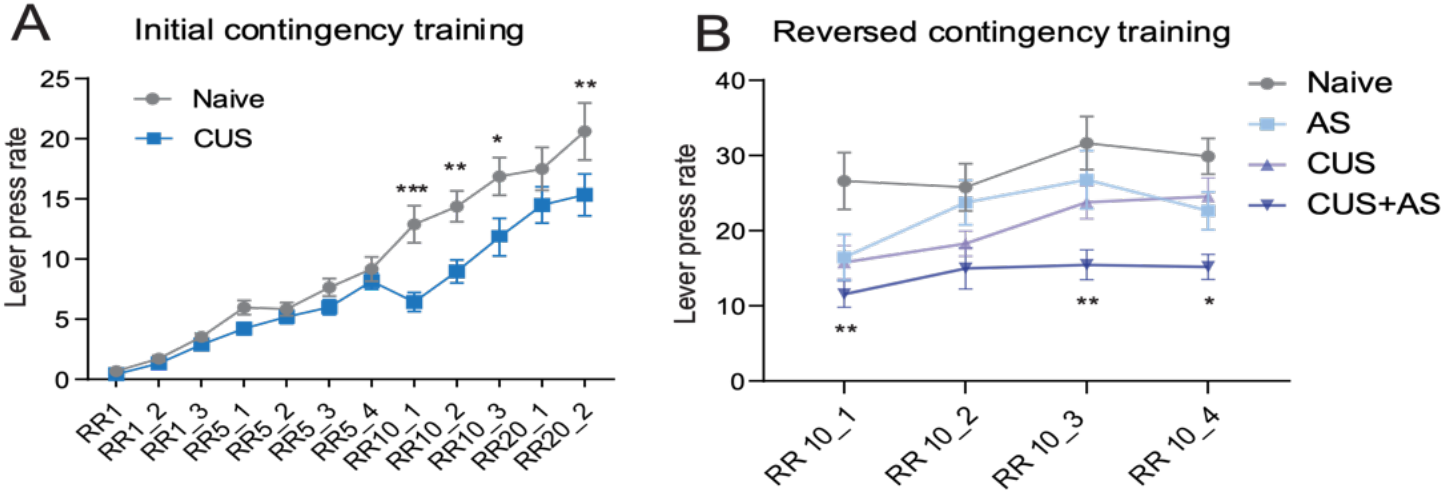
Contingency Learning Following Chronic and Acute Stress Exposure (adapted from Mor et al., 2022) **A**) Group mean lever pressing rates during initial learning. A repeated measures ANOVA showed pressing rates increased over the training, F_(11,52)_ = 30.79, *p*<.001, but that this increase differed between the CUS and Naïve rats yielding a stress × training interaction, F_(11,52)_=2.29, *p*=.023. Pressing rates were significantly lower for CUS than Naïve rats, F_(1,62)_ = 6.75, p =.012, suggesting attenuated rigor after CUS. **B**) Group mean lever pressing rates during reversal learning. A repeated measures ANOVA revealed that pressing rates increased significantly over sessions, *F*_(3,165)_=15.96, *p*<.001, and there was no interaction was between group and sessions, *F*_(9,165)_=1.81, *p*= .070. There was, however, evidence of a difference in lever pressing between treatment groups *F*_(3,55)_=6.55, *p*< .001, with main effects for CUS *F*_(1,55)_=12.38, *p*<.001 as well as AS *F*_(1,55)_=7.42, *p*=.008. Bonferroni adjusted pairwise comparisons indicated reduced pressing rates in *CUS with AS* when compared with naïve rats on training days one (*p*= .009), two (*p*= .002) and four (*p*=.012). Bars represent means ±SEM. \**p*<.05, \**p*<.01, \*\**p*<.001, *** *p*<.0001.

**Supplementary figure 2.**
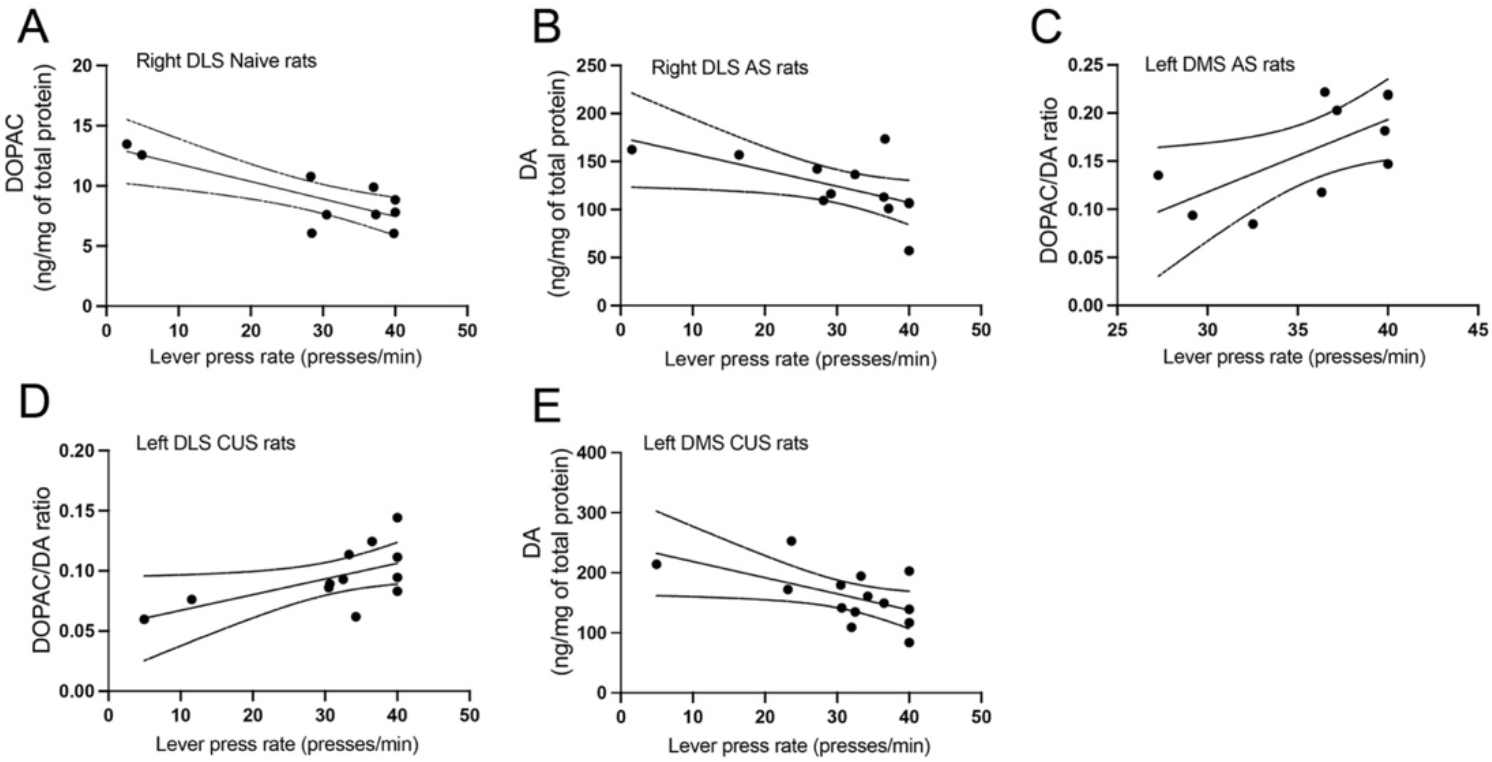
Correlations between monoamine levels and behavioral performance during reversal training. During reversal training, pressing rates in both naïve and AS rats decreased with increased dopaminergic activity in the right DLS, with negative correlations with DOPAC in naïve (r^2^=.627, *p*=.006) (A) and DA in AS rats (r^2^=.411, *p*=.024) (**B**). In the left DMS, AS and CUS led to opposite associations, with pressing increasing in AS with increased DOPAC/DA ratios (r^2^=.441, *p*=.0360) (**C**) and decreasing with increased DA in CUS rats (r^2^=.313, *p*=.037) (**E**). In contrast to what was found in the left DMS though, CUS rats pressing increased with increased DOPAC/DA in the left DLS (r^2^=.355, *p*=.04) (**D**).

**Supplementary figure 3.**
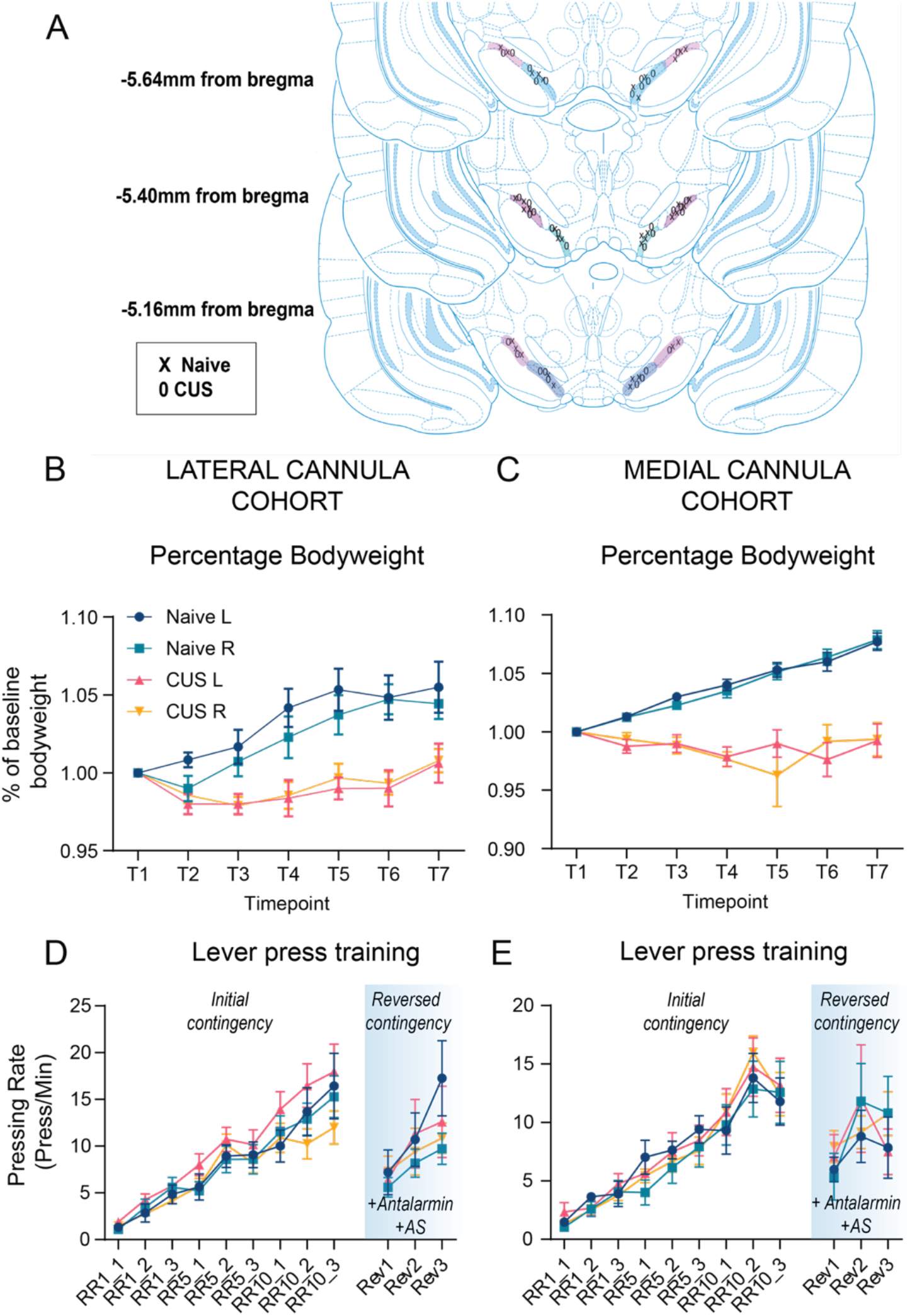
The effect of lateralized CRFR1-antagonist infusions on body weight and instrumental training. **A**) Map of cannula placements for lateral (in pink) and medial (in blue) cohorts. Percentage bodyweight for the lateral cannula cohort (**B**) and the medial cannula cohort (**C**) across 2 weeks of CUS or handling. Bodyweight was collected every second day. 2×7 ANOVA revealed a significant time x stress interaction, in both the lateral SNpc cohort: F_(2.12, 67.86)_=15.34, p<.001 and the medial SNpc cohort: F_(3.26, 91.20)_=3.26, p<.001. **D-E**) Lever press training for lateral (D) and medial (E) cannula animals during the initial and reversed contingency training. Throughout training, a 2×2×9 mixed ANOVA showed that all groups quickly acquired lever contingencies, as pressing rates increased significantly over the training sessions. Both medial and lateral cannula cohorts showed an overall effect of training F_(2.71, 70.37)_ = 65.07, *p*<.001 and F_(2.93, 87.86)_ = 67.72, *p*<.001, respectively, with no significant differences between group Naïve and CUS (all Fs<1) or implantation location, all Fs<2, *p*> .05. A mixed 2×2×3 ANOVA revealed a significant main effect as pressing rate increased over sessions: medial cohort *F*_(2, 60)_=6.87, *p*=.002, lateral cohort *F*_(2, 52)_=20.25, *p*<.001. No interactions between session x stressor was found for either cohort: medial infusion *F*_(1, 30)_=1.15, *p*=.293, lateral infusions *F*_(2, 52)_=1.18, *p*=.317, neither any interaction between session and infusion side (left or right): medial infusion *F*_(1, 30)_=2.463, *p*=.127, lateral infusions *F*_(2, 52)_=34.40, *p*=.081. Bars represent means ±SEM. \**p*<.05, \**p*<.01, \*\**p*<.001, *** *p*<.0001.

**Table S1:** Tests of within-subjects contrasts and between subject effects for each of the monoamines, metabolites and utilisation rates measured in DMS and DLS.

|  | DMS |  |  |  |  |  |
| --- | --- | --- | --- | --- | --- | --- |
|  | Main effect for side |  | Main effect for AS |  | Main effect for CUS |  |
| DA | F (1, 44) = 1.315 | P=.258 | **F (1,44) = 4.174** | **P=.047** | F (1,44) = 1.311 | P=.258 |
| DOPAC | F (1,44) = 3.352 | P=.074 | F (1,44) = .079 | P=.78 | F (1,44) = 0 | P=.989 |
| DOPAC/DA | F (1,41) = .833 | P=.367 | **F (1,41) = 9.665** | **P=.003** | **F (1,41) = 4.295** | **P=.045** |
|  | DLS |  |  |  |  |  |
|  | Main effect for side |  | Main effect for AS |  | Main effect for CUS |  |
| DA | F (1,40) = .366 | P=.549 | F (1,40) = 1.269 | P=.267 | F (1,40) = .242 | P=.626 |
| DOPAC | F (1,40) = 2.141 | P=.151 | F (1,40) = .195 | P=.661 | F (1,40) = 3.048 | P=.089 |
| DOPAC/DA | F (1,44) = .96 | P=.758 | **F (1,44) = 8.637** | **P=.005** | F (1,44) = 2.381 | P=.13 |

|  | DMS |  |  |  |  |  |  |  |
| --- | --- | --- | --- | --- | --- | --- | --- | --- |
|  | Side*AS*CUS Interaction |  | side*AS interaction |  | side*CUS interaction |  | AS*CUS interaction |  |
| **DA** | **F(1,44) = 4.09** | **P=.049** | **F(1,44) = 5.468** | **P=.024** | **F(1,44) = 4.049** | **P=0.05** | F(1,44) = .03 | P.862 |
| DOPAC | **F(1,44) = 5.173** | **P=.028** | F(1,44) = 13.357 | P=.371 | F(1,44) = 2.256 | P=0.14 | **F(1,44) = 10.047** | **P=0.003** |
| DOPAC/DA | F(1,41) = .624 | P=.434 | **F(1,41) = 8.561** | **P=.006** | F(1,41) = 2.723 | P=0.107 | **F(1,41) = 20.534** | **P<.000** |
|  | DLS |  |  |  |  |  |  |  |
|  | Side*AS*CUS Interaction |  | side*AS interaction |  | side*CUS interaction |  | AS*CUS interaction |  |
| DA | F (1,40) = .622 | P=.435 | F(1,40) = 2.058 | P=.159 | F(1,40) = .104 | P=.749 | **F(1,40) = 8.123** | **P=.007** |
| DOPAC | F(1,40) = .226 | P=.637 | F(1,40) = .572 | P=.545 | F(1,40) = 1.103 | P=.3 | **F(1,40) = 14.942** | **P<.000** |
| DOPAC/DA | F(1,44) = .096 | P=.758 | **F(1,44) = 4.68** | **P=.036** | F(1,44) = .661 | P=.42 | F(1,44) = 1.956 | P=.169 |

